# Mendelian randomization: a premature burial?

**DOI:** 10.1101/021386

**Authors:** George Davey Smith

## Abstract

Mendelian randomization is a promising approach to help improve causal inference in observational studies, with widespread potential applications, including to prioritization of pharmacotherapeutic targets for evaluation in RCTs. From its initial proposal the limitations of Mendelian randomization approaches have been widely recognised and discussed, and recently Pickrell has reiterated these^1^. However this critique did not acknowledge recent developments in both methodological and empirical research, nor did it recognise many future opportunities for application of the Mendelian randomization approach. These issues are briefly reviewed here.

---

Whilst providing an appropriate note of caution with respect to the interpretation of Mendelian randomization study findings^1^, Pickrell’s reasons for viewing Mendelian randomization as having so far failed to fulfil its promise are based on some misconceptions.

## 30 years of Mendelian randomization?

Pickrell says that the “first reason for scepticism is that in the nearly 30 years of Mendelian randomization, arguably no new causal relationship has been identified with this approach and subsequently verified in a randomized controlled trial” 1. The thirty year figure appears to have derived from the notion that the method began to be applied following a publication by Martijn Katan in 1986^2^. Katan’s insightful contribution was a letter to the Lancet which proposed the use of an *APOE* genetic marker for cholesterol level to interrogate the claim that low circulating cholesterol increases the risk of cancer. The major issue with observational studies that had examined the association between circulating cholesterol level and cancer was that early stages of disease could lower cholesterol level, but would not influence genotype (i.e. the reverse causation problem that exists for conventional observational studies would not influence a genetic association). Katan also pointed out that genotypes would be related to long-term (since birth) differences in cholesterol levels. This would provide a powerful test of the low cholesterol-cancer risk hypothesis. No data were presented in the letter, and it did not use the term Mendelian randomization. In the subsequent 17 years the letter was only cited twice^3 4^, neither time in relation to its proposed methodology. No “Mendelian randomization” studies followed its publication, and it has only become widely cited (one suspects often by people who have not read it, given the disconnect between what the citations suggest was in the letter and what was actually there) after it was referenced as one of the antecedents of Mendelian randomization in the first extended presentation of the approach in 2003^5^, and then we reprinted it in the *International Journal of Epidemiology* in 2004^6 7^

Pickrell is not alone in misstating the time span over which Mendelian randomization studies have been performed, with others claiming that the method has been used in epidemiological studies for more than 20 years^8^. Some confusion is created by the fact that the term “Mendelian randomization” was introduced in a very different context, in studies utilizing whether or not acute myeloid leukemia patients had HLA compatible siblings as a way of obtaining evidence as to the survival benefits of bone marrow transplants.^9^ Strangely, this paper is quoted as the source of study designs that use “genetically determined differences in exposure to test whether a biomarker affects disease”, and to provide evidence of the more than two decades over which the use of “genetic variation in a biomarker to deduce the causal effects of that biomarker on a disease” has been in use^10^. In fact the term Mendelian randomization has only been used in its current sense since the early 2000s^5 11 12 13^.

This archaeology of the concept of Mendelian randomization (provided in more depth elsewhere)^14^ is of relevance to Pickrell’s critique, as the main plank of his first “reason for scepticism” is based on a misperception of time scale over which such studies have been carried out. Indeed it is only since the era of robust identification of common genetic variant associations with phenotypes in the post-GWAS era that Mendelian randomization studies have generally become feasible, reflected in the very considerable increase in publication of Mendelian randomization studies over the last five years.

## No evidence of Mendelian randomization studies leading to clinical trials?

Given the well documented long time-course required for drug development^15^, or the development of non-drug public health and clinical interventions, it is of course entirely unsurprising that there has not been a cascade of new causal relationships that have been identified with this approach and subsequently verified in randomized controlled trials, as Pickrell seems to expect should have occurred in the 12 years since the use of this study design was first articulated. He suggests, indeed, that there are no such cases. This is incorrect – remarkably (given the short time period over which Mendelian randomization studies have been applied, and the time required for drug, public health and clinical intervention development through to completion of long-term phase 3 trials) – there are several cases. For example, proprotein convertase subtilisin/kexin type 9 serine protease (*PCSK9)* genetic variation was identified as relating to LDL cholesterol level and coronary heart disease in 2006^16^ and the two largest randomised controlled trials (RCTs) of monoclonal antibodies targeting PCSK9 that have appeared since publication of that Mendelian randomization study, both suggesting ^17 18^ a reduction in cardiovascular events, as does a meta-analysis including smaller trials ^19^.

In conventional observational studies lipoprotein-associated phospholipase A2 (L_p_-PLA_2_) levels have been shown to predict coronary heart disease (CHD) risk for many years, with this being apparently independent of conventional CHD risk factors (see e.g. the 2010 large scale overview of such studies^20^). This led to the development of pharmacotherapeutic agents which lowered L_p_-PLA_2_. However genetic studies on the V279F variant in *PLA2G7*, which is common in East Asian but not European-origin populations and is associated with particularly low levels of L_p_-PLA_2_, suggested that there would not be a substantial effect of lowering L_P_-PLA_2_ levels on CHD risk (see e.g. a 2010 metaanalysis of these)^21^. Subsequently large scale trials have failed to find the benefit that was anticipated from a naive interpretation of the observational epidemiological data^22 23 24^. Whilst other trials are ongoing the magnitude of anticipated benefit (if any) has certainly been down-scaled as a result of the Mendelian randomization studies.

Additionally there are cases where genetic studies and the development of therapies co-evolved. For example, findings with respect to Niemann-Pick C1-Like 1 Protein (NPC1L1) genetic variation and ezetimibe, the drug that targets NPC1L1, were cited in a recent review of Mendelian randomization applied to drug development as being apparently discrepant^25^. Recent large-scale genetic and RCT data now suggest the findings are concordant ^26 27 28^. Furthermore there are several examples of ongoing trials testing expectations from Mendelian randomization studies, for example trials of IL6 receptor blockade^29^ that will evaluate the predictions from Mendelian randomization studies^30 31^.

As well as contributing to the elucidation of some drug targets, Mendelian randomization has also deprioritized the development of some pharmacotherapeutic agents. For example the further development of therapies targeting C-reactive protein (CRP) levels was not encouraged by the many Mendelian randomization studies suggesting CRP was not causal with respect to a range of cardiometabolic outcomes ^32 33 34 35^, and targeting the endothelial lipase gene *(LIPG* Asn396Ser) to lower HDL cholesterol was discouraged by a large-scale Mendelian randomization study of this.^36^

To support his argument Pickrell provides a supplementary table presenting a comparison of Mendelian randomization studies and RCTs supposedly investigating the same issue, implying that this is a comprehensive list of Mendelian randomization studies. But in fact this includes a very limited number of studies included purely as illustrative examples in two overviews of Mendelian randomization. But there are a very large number of Mendelian randomization studies – for a partial but more systematic listing, see Boef et al^37^. This presents nearly 200 studies, but remains only a sample since it is based on search and selection criteria that have missed a large number of such studies (e.g. consortia papers, which are increasingly common in the field, those using different terminologies but nonetheless applying Mendelian randomization methods, etc.). An attempt to evaluate Mendelian randomization on the basis of a (in any case flawed) comparison should attempt a somewhat more systematic approach. Furthermore Pickrell fails to recognise the widely understood limitations of RCTs. As just one example in the table he suggests there are no RCTs of alcohol reduction and blood pressure. This is far from the truth, already by 2001 a widely cited metaanalysis included 15 such RCTs^38^, and more have appeared since. However such trials have problems in producing large sustained changes in alcohol consumption. The power of Mendelian randomization to provide useful evidence in situations where trials are difficult or impossible to successfully implement is a very considerable strength, not a weakness, of the approach.

## Pleiotropy: what’s new?

The second reason for scepticism that Pickrell raises, that of pleiotropy, has been discussed in very considerable detail, from the initial 2003 paper5 onwards, e.g. recently by Glymour et al^39^ and Vanderweele^40^. As we will see, the simulation he provides does not add greatly to what is known and widely recognised about the potential effect of pleiotropy. Furthermore the paper fails to acknowledge the extensive (and empirically useful) work in the instrumental variables field, much of which has been directly applied to and utilised in the Mendelian randomization context, and is well known to practitioners of IV analysis. These methods allow for robustness checks, and estimation which remains valid under relaxed IV assumptions. For a limited sample of this literature, see, e.g,^41 42 43 44 45 46 47 48^.

Against the background of this exciting work Pickrell makes two unreliable observations. First, he cites Bulik-Sullivan’s groundbreaking work using whole-genome LD regression approaches^49^ as showing that “genetic variants that influence HDL cholesterol levels have correlated effects on whether an individual went to college. A naïve interpretation of this might suggest the (rather nonsensical) conclusion that HDL-raising drugs should increase education levels”.^1^ This makes it sound as though Bulik-Sullivan’s work is an example of Mendelian randomization, which it is not - it simply suggests there is a genetic correlation, that could be generated by vertical (real) phenotypic effects, or be through horizontal (spurious) pleiotropy, or a combination of the two. Furthermore there is no direction implied by there being a genetic correlation, whilst the nonsensical conclusion that Pickrell highlights assumes that direction of effect can be inferred. This is not the case. It is the case, however, that higher college education - and things that follow on from this, like greater awareness of healthy diet, improved exercise behaviour, the use of medications etc - will influence HDL cholesterol. The genetic correlation could, at least in part, be through phenotypic effects.

Second, Pickrell states that “it is sometimes suggested that using a large number of genetic variants in Mendelian randomization (combined into a single score) offers a way to partially avoid this problem”^1^, and the simulations that are then performed examine this situation. Unsurprisingly, if you simulate a situation with pleiotropy you demonstrate the existence of pleiotropic effects. However approaches to utilising multiple genetic variants in Mendelian randomization studies have considered comparing potentially large numbers of independent estimates, which allow evaluation of the extent to which pleiotropy may be biasing effect estimates, as the particular strength of this situation. They do not suggest that combinations into a single score provide such reassurance. For example, in an introductory paper on Mendelian randomization published some years ago, it was suggested that

> “In some cases, it may be possible to identify two separate genetic variants, which are not in linkage disequilibrium with each other, but which both serve as proxies for the environmentally modifiable risk factor of interest. If both variants are related to the outcome of interest and point to the same underlying association, then it becomes much less plausible that reintroduced confounding explains the association, since it would have to be acting in the same way for these two unlinked variants. This can be likened to RCTs of different blood pressure-lowering agents, which work through different mechanisms and have different potential side effects. If the different agents produce the same reductions in cardiovascular disease risk, then it is unlikely that this is through agent-specific (pleiotropic) effects of the drugs; rather, it points to blood pressure lowering as being key. The latter is indeed what is in general observed^50^. In another context, two distinct genetic variants acting as instruments for higher body fat content have been used to demonstrate that greater adiposity is related to higher bone mineral density^51^. With the large number of genetic variants that are being identified in genome wide association studies in relation to particular phenotypes—e.g. >50 independent variants that are related to height; >90 that are related to total cholesterol and >20 related to fasting glucose—it is possible to generate many independent combinations of such variants and from these many independent instrumental variable estimates of the causal associations between an environmentally modifiable risk factor and a disease outcome. The independent estimates will not be plausibly influenced by any common pleiotropy or LD-induced confounding, and therefore if they display consistency this provides strong evidence against the notion that reintroduced confounding is generating the associations”^52^.

A simple use of multiple genetic instruments for evaluating the plausibility of distortion of Mendelian randomization findings by pleiotropy is illustrated in the figure. It is increasingly improbable for 2, 3, 4 or more genetic variants out of LD with each other to lead to precisely the same quantitative causal effect estimate due to perfectly balancing pleiotropy. Here we see data from 9 SNPs from 6 genes which lead to remarkably similar predicted causal effects of LDL cholesterol on CHD risk.

The more recent developments in instrumental variables approaches referenced earlier move well beyond even this interrogation of the potential influence of pleiotropy in the Mendelian randomization setting, and provide an extensive range of sensitivity analyses that can inform interpretation of the findings^41 42 43 47^. Ironically, one of the conclusions of Pickrell’s exciting recent work on shared genetic influences on human traits^53^ - that “the effect sizes of the variants on the different traits appear to be largely uncorrelated”^53^ – provides the basis for one approach to utilising multiple potentially pleiotropic genetic variants in the Mendelian randomization context.^43^

## Fulfilling the potential of Mendelian randomization?

The suggestions Pickrell makes regarding fulfilling the promises of Mendelian randomization are useful considerations. However they fail to anticipate some transformational levels of evidence that can be provided by Mendelian randomization approaches. To give a short selection of these related to just one issue, that of pharmaceutical target evaluation:

(1) Predicting the comprehensive phenotypic effect of pharmaceutical treatments (including unwanted side effects) based on interrogation of the phenome-wide associations of a genetic proxy. One example suggests that IL1 manipulation – currently being trialled in autoimmune disease contexts – may increase rather than decrease cardiovascular risk.^54^

(2) The separation of on-target from off-target effects of pharmacotherapy. A recent example suggests that the increase in diabetes risk seen in statin trials is a consequence of its mechanism of reducing cholesterol, rather than an off-target side effect unrelated to the pathway thought which cholesterol is reduced^55^

(3) The separation of specific mechanism from intermediate phenotype effects. In the above statin example is the on-target effect could be due to HMGCoA reductase manipulation or to cholesterol lowering, in which case all approaches to cholesterol reduction would have the same influence on diabetes risk, proportional to their success in actually reducing cholesterol level

(4) Providing evidence on generalizability of findings from RCTs. It is not feasible to conduct adequately powered large scale RCTs of a treatment in every possible subgroup of the global population (defined by combinations of age, gender, ethnicity, an extensive range of comorbidities, for example), but Mendelian randomization studies using genetic proxies could be carried out at a tiny fraction of the cost

(5) Providing evidence on combined drug treatment – answering the question ‘will combined therapy produce additional benefits?’ – through analysis of combination of genetic variants mimicking the different drugs, in the factorial Mendelian randomization design^28^.

Recent commentaries, illustrating the considerable potential in this field, are available elsewhere.^56^ ^57^. Similar and greatly expanded lists could be created in many other domains of research, translation and practice.

## Mendelian randomization in its place

Pickrell draws attention to some limitations of Mendelian randomization, and many others of course exist and have been discussed at length in the published literature. Ways of moving beyond these are being developed, however. There are some fundamental issues not discussed by Pickrell that deserve attention as Mendelian randomisation develops. First, most of the currently available GWAS data are not the ones that are required to answer some of the most important questions in medicine and related disciplines. Mendelian randomization has largely been applied to disease aetiology (e.g. in case-control studies of particular diseases), but this does not have any direct bearing on therapy. For example, GWAS clearly identified a proxy for smoking intensity as the strongest common genetic variant related to lung cancer, reaffirming the causal influence of smoking on lung cancer. However, once lung cancer has developed stopping smoking is not an effective treatment. It is likely that many – diseases demonstrate a similar disconnect between triggers (which cannot be reversed through the same process once the disease has been initiated) and factors related to progression and prognosis. Mendelian randomization studies of disease incidence are a powerful tool for identifying applied to studies of disease progression offers to provide evidence regarding factors that could prevent disease – such as smoking and lung cancer as a proof of principle example) for some conditions – e.g. CHD – factors that relate to disease incidence (higher blood pressure, higher cholesterol, and smoking) relate to disease progression and future events. However in other conditions this will not be the case. Mendelian randomization applied to studies of disease progression may provide evidence regarding factors that could be manipulated to improve prognosis, and establishing such studies would represent a major advance.

Second, Mendelian randomization generally utilises genetic variants which influence life-long exposure to different levels of a potential risk factor. This means that such studies can establish the very long-term effects of such exposures^58^, and has advantages in terms of demonstrating what would be possible with preventative initiatives starting in early life, and the public health effect of factors that produce population-level shifts in a risk factor such as blood pressure or cholesterol. However a potential downside is that if exposure at one period of life leads to a change in risk that is not reversible, the Mendelian randomization findings would not be reproduced in a trial modifying the factor in later life. For example, if lower levels of antioxidant exposure (e.g. vitamin C, uric acid, bilirubin) in infancy or childhood lead to changes in the arterial wall which cannot be reversed by later antioxidant supplementation, Mendelian randomization findings would correctly suggest adverse effects of low antioxidant levels, but this would not translate into benefit of antioxidant treatment. In principal it could be possible to utilise gene by environment interactions to identify critical periods during which exposures act^52^, but in reality obtaining datasets for this of adequate sample size, to establish robust gene by environment interactions, will be a seriously limiting factor. This illustrates the need to combine Mendelian randomisation evidence with other sources of information

As indicated above, Mendelian randomization provides one plank of evidence on a particular question. The evaluation of any particular question requires the triangulation of evidence from different approaches, particularly from approaches which, whilst potentially biased, suffer from nonassociated forms of bias^59 60 61^. The approach provides a potentially powerful narrowing down of hypotheses for further testing, for example, the selection of candidate pharmacotherapeutic agents for evaluation in expensive RCTs. As we said in the initial extended presentation of Mendelian randomization “It is probably fair to say that the method offers a more robust approach to understanding the effect of some modifiable exposures on health outcomes than does much conventional observational epidemiology. Where possible randomized controlled trials remain the final arbiter of the effects of interventions intended to influence health, however”^5^.

**Figure.**
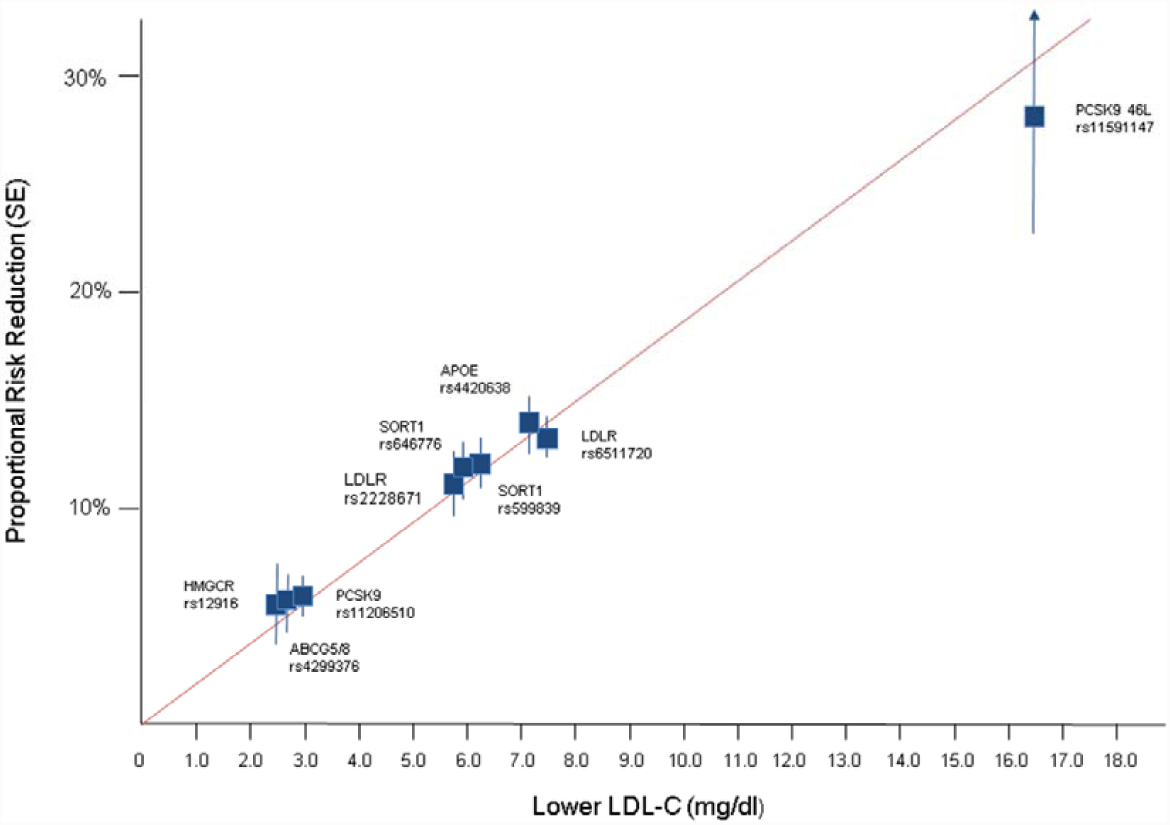
**Effect of lower LDL-C on risk of CHD** [adapted from Ference et al. (2012)^i^]. Boxes represent the proportional risk reduction (1-OR) of CH D for each exposure allele plotted against the absolute magnitude of lower LDL-C associated with that allele (measured in mg/dl).SNPs are plotted in order of increasing absolute magnitude of associations with lower LDL-C. The line (forced to pass through the origin) represents the inc rease in proportional risk reduction of CHD per unit lower long-term exposure to LDL-C.

i Ference BA, Yoo W, Alesh I, Mahajan N, Mirowska KK, Mewada A, Kahn J, Afonso L, Williams KA Sr, Flack JM. Effect of long-term exposure to lower lowdensity lipoprotein cholesterol beginning early in life on the risk of coronary heart disease: a Mendelian randomization analysis. J. Am. Coll. Cardiol 2012, 60, 2631–2639

## Acknowledgements

Thanks to Stephen Burgess, Dave Evans, Marcus Munafo, Caroline Relton and Nic Timpson for comments on an earlier draft of this piece.

## References

1 Pickrell J. Fulfilling the promise of Mendelian randomization. BioRxiv 2015; doi: http://dx.doi.org/10.1101/018150.

2 Katan M. Apolipoprotein E isoforms, serum cholesterol, and cancer. Lancet 1986;1:507–8.

3 Kritchevsky SB, Kritchevsky D. Serum cholesterol and cancer risk: an epidemiologic perspective. Annu Rev Nutr. 1992;12:391–416.

4 Davignon J, Gregg RE, Sing CF. Apolipoprotein E polymorphism and atherosclerosis. Arteriosclerosis. 1988;8:1–21.

5 Davey Smith G, Ebrahim S. 'Mendelian randomization': can genetic epidemiology contribute to understanding environmental determinants of disease? Int J Epidemiol.2003;32:1–22.

6 Katan M. Apolipoprotein E isoforms, serum cholesterol, and cancer. Int J Epidemiol 2004;33:9

7 Katan M. Commentary: Mendelian randomization 18 years on. Int J Epidemiol 2004;33:10–11

8 Freeman G, Cowling BJ, Schooling CM. Power and sample size calculations for Mendelian randomization studies using one genetic instrument. Int J Epidemiol 2013;42:1157–1163.

9 Gray R, Wheatley K. How to avoid bias when comparing bone marrow transplantation with chemotherapy. Bone Marrow Transplant 1991;7:9–12.

10 Schooling CM,Freeman G, Cowling BJ. Mendelian randomization and estimation of treatment efficacy for chronic diseases. Am J Epidemiol 2013;177:1128–33.

11 Youngman LD, Keavney BD, Palmer A et al. Plasma fibrinogen and fibrinogen genotypes in 4685 cases of myocardial infarction and in 6002 controls: test of causality by 'Mendelian randomization’. Circulation 2000;102(Suppl. II):31–32.

12 Clayton D, McKeigue PM. Epidemiological methods for studying genes and environmental factors in complex diseases. Lancet 2001; 358:1356–60.

13 Fallon UB, Ben-Shlomo Y, Davey Smith G. Homocysteine and coronary heart disease. Heart Online 14 March 2001

14 Davey Smith G. Commentary. Capitalizing on Mendelian randomization to assess the effects of treatments. Journal of the Royal Society of Medicine 2007;100:432–435 (also at http://www.jameslindlibrary.org/illustrating/articles/capitalising-on-mendelian-randomization-to-assess-the-effects-of)

15 David Cook, Dearg Brown, Robert Alexander, Ruth March, Paul Morgan, Gemma Satterthwaite, Menelas N. Pangalos. Lessons learned from the fate of AstraZeneca's drug pipeline: a five-dimensional framework. Nature Drug Discovery 2014; 13: 419–431.

16 Cohen JC, Boerwinkle E, Mosley TH Jr, Hobbs HH. Sequence variations in PCSK9, low LDL, and protection against coronary heart disease. N Engl J Med 2006;354:1264–72.

17 Sabatine MS, Giugliano RP, Wiviott SD, Raal FJ, Blom DJ, Robinson J, Ballantyne CM, Somaratne R, Legg J, Wasserman SM, Scott R, Koren MJ, Stein EA; Open-Label Study of Long-Term Evaluation against LDL Cholesterol (OSLER) Investigators. Efficacy and safety of evolocumab in reducing lipids and cardiovascular events. N Engl J Med 2015;372:1500–9.

18 Robinson JG, Farnier M, Krempf M, Bergeron J, Luc G, Averna M, Stroes ES, Langslet G, Raal FJ, El Shahawy M, Koren MJ, Lepor NE, Lorenzato C, Pordy R, Chaudhari U, Kastelein JJ; ODYSSEY LONG TERM Investigators. Efficacy and safety of alirocumab in reducing lipids and cardiovascular events. N Engl J Med. 2015 Apr 16; 372(16):1489–99. doi: 10.1056/NEJMoa1501031. Epub 2015 Mar 15.

19 Navarese EP, Kolodziejczak M, Schulze V, Gurbel PA, Tantry U, Lin Y, Brockmeyer M, Kandzari DE, Kubica JM, D'Agostino RB Sr, Kubica J, Volpe M, Agewall S, Kereiakes DJ, Kelm M. Effects of Proprotein Convertase Subtilisin/Kexin Type 9 Antibodies in Adults With Hypercholesterolemia: A Systematic Review and Meta-analysis. Ann Intern Med 2015; doi: 10.7326/M14-2957.

20 The Lp-PLA Studies Collaboration. Lipoprotein-associated phospholipase A2 and risk of coronary disease, stroke, and mortality: collaborative analysis of 32 prospectives studies. Lancet 2010;375:1536–1544.

21 Wang Q, Hao Y, Mo X, Wang L, Lu X, Huang J, Cao J, Li H, Gu D. PLA2G7 gene polymorphisms and coronary heart disease risk: a meta-analysis. Thrombosis Research 2010;126:498–503.

22 STABILITY Investigators, White HD, Held C, Stewart R, Tarka E, Brown R, Davies RY, Budaj A, Harrington RA, Steg PG, Ardissino D, Armstrong PW, Avezum A, Aylward PE, Bryce A, Chen H, Chen MF, Corbalan R, Dalby AJ, Danchin N, De Winter RJ, Denchev S, Diaz R, Elisaf M, Flather MD, Goudev AR, Granger CB, Grinfeld L, Hochman JS, Husted S, Kim HS, Koenig W, Linhart A, Lonn E, López-Sendón J, Manolis AJ, Mohler ER 3rd, Nicolau JC, Pais P, Parkhomenko A, Pedersen TR, Pella D, Ramos-Corrales MA, Ruda M, Sereg M, Siddique S, Sinnaeve P, Smith P, Sritara P, Swart HP, Sy RG, Teramoto T, Tse HF, Watson D, Weaver WD, Weiss R, Viigimaa M, Vinereanu D, Zhu J, Cannon CP, Wallentin L. Darapladib for preventing ischemic events in stable coronary heart disease. N Engl J Med 2014;370:1702–11.

23 Nicholls SJ, Kastelein JJ, Schwartz GG, Bash D, Rosenson RS, Cavender MA, Brennan DM, Koenig W, Jukema JW, Nambi V, Wright RS, Menon V, Lincoff AM, Nissen SE; VISTA-16 Investigators. Varespladib and cardiovascular events in patients with an acute coronary syndrome: the VISTA-16 randomized clinical trial. JAMA 2014;311:252–62.

24 O’Donoghue ML, Braunwald E, White HD, Steen DL, Lukas MA, Tarka E, Steg PG, Hochman JS, Bode C, Maggioni AP, Im K, Shannon JB, Davies RY, Murphy SA, Crugnale SE, Wiviott SD, Bonaca MP, Watson DF, Douglas Weaver W, Serruys PW, Cannon CP, for the SOLID-TIMI 52 Investigators. Effect of Darapladib on Major Coronary Events After an Acute Coronary Syndrome. The SOLID-TIMI 52 Randomized Clinical Trial. JAMA 2014;312:1006–1015.

25 Mokry LE, Ahmad O, Forgetta V, Thanassoulis G, Richards JB. Mendelian randomisation applied to drug development in cardiovascular disease: a review. J Med Genet. 2015;52:71–9.

26 Cannon CP, Blazing MA, Giugliano RP, McCagg A, White JA, Theroux P et al. Ezetimibe Added to Statin Therapy after Acute Coronary Syndromes. New Eng J Med 2015;372:2387–2397.

27 The Myocardial Infarction Genetics Consortium Investigators. Inactivating Mutations in NPC1L1 and Protection from Coronary Heart Disease. N Engl J Med 2014;371:2072–2082.

28 Ference BA, Yoo W, Alesh I, Mahajan N, Mirowska KK, Mewada A, Kahn J, Afonso L, Williams KA Sr, Flack JM. Effect of long-term exposure to lower low-density lipoprotein cholesterol beginning early in life on the risk of coronary heart disease: a Mendelian randomization analysis. J. Am. Coll. Cardiol 2012, 60, 2631–2639

29 Ridker PM, Lüscher TF. Anti-inflammatory therapies for cardiovascular disease. Eur Heart J. 2014;35:1782–91.

30 IL6R Genetics Consortium Emerging Risk Factors Collaboration. Interleukin-6 receptor pathways in coronary heart disease: a collaborative meta-analysis of 82 studies. Lancet 2012;379:1205–1213.

31 The Interleukin-6 Receptor Mendelian Randomisation Analysis (IL6R MR) Consortium. The interleukin-6 receptor as a target for prevention of coronary heart disease: a mendelian randomisation analysis. The Lancet 2012;379:1214–1224.

32 Timpson NJ, Lawlor DA, Harbord RM, Gaunt TR, Day INM, Palmer LJ, Hattersley AT, Ebrahim S, Lowe GDO, Rumley A, Davey Smith G. C-reactive protein and its role in metabolic syndrome: Mendelian randomisation study. Lancet 2005;366:1954–1959.

33 Zacho J, Tybjærg-Hansen A, Jensen JS, Grande P, Sillesen H, Nordestgaard BG. Genetically Elevated C-Reactive Protein and Ischemic Vascular Disease. N Engl J Med 2008;359:1897–1908.

34 Lawlor DA, Harbord RM, Timpson NJ, Lowe GDO, Rumley A, Gaunt TR, Baker I, Yarnell JWG, Kivimäki M, Kumari M, Norman PE, Jamrozik K, Hankey GJ, Almeida OP, Flicker L, Warrington N, Marmot MG, Ben-Shlomo Y, Palmer LJ, Day INM, Ebrahim S, Davey Smith G. The association of C-reactive protein and CRP genotype with coronary heart disease: Findings from five studies with 4,610 cases amongst 18,637 participants. PLoS One 2009;8:e3011.

35 Wensley F, Gao P, Burgess S et al, on behalf of the C-Reactive Protein Coronary Heart Disease Genetics Collaboration (CCGC). Association between C reactive protein and coronary heart disease: mendelian randomisation analysis based on individual participant data. BMJ 2011;342:d548.

36 Voight B, Peloso GM, Orho-Melander et al. Plasma HDL cholesterol and risk of myocardial infarction: a mendelian randomisation study. Lancet 2012;380:572–580.

37 Boef AGC, Dekkers OM, le Cessie S. Mendelian randomization studies: a review of the approaches used and the quality of reporting. International Journal of Epidemiology 2015; doi:10.1093/ije/dyv071.

38 Xin X, He, J, Frontini MG, Ogden LG, Motsamai OI, Whelton PK. Effects of alcohol consumption on blood pressure: a meta-analysis of randomised controlled trials. Hypertension 2001; 38:1112–7

39 Glymour MM, Tchetgen Tchetgen EJ, Robins JM. Credible Mendelian randomization studies: approaches for evaluating the instrumental variable assumptions. Am J Epidemiol 2012;175:332–9.

40 VanderWeele TJ, Tchetgen Tchetgen EJ, Cornelis M, Kraft P. Methodological challenges in mendelian randomization. Epidemiology. 2014;25:427–35.

41 Kang H, Zhang A, T. Cai TT, Small DS. Instrumental Variables Estimation with Some Invalid Instruments and its Application to Mendelian Randomization. Journal of the American Statistical Association 2015; DOI: 10.1080/01621459.2014.994705

42 Kang H, Cal TT, Small SD. Robust confidence intervals with possibly invalid instruments. arXiv:1504.03718

43 Bowden J, Burgess S, Davey Smith G. Mendelian randomization with invalid instruments: effect estimation and bias detection through Egger regression. Int J Epidemiol 2015:44:512–525.

44 Slichter D. Testing Instrument Validity and Identification with Invalid Instruments. SOLE 2014; http://www.sole-jole.org/14436.pdf.

45 Kolesar M, Chetty R, Friedman J, Glaeser E, Imbens G. Identification and Inference with Many Invalid Instruments. Journal of Business & Economic Statistics 2014: DOI: 10.1080/07350015.2014.978175

46 Han C. Detecting Invalid Instruments Using L1-GMM. Economic Letters 2008;101:285–287.

47 Smith JG, Luk K, Schulz C, Engert JC, Do R, Hindy G, Rukh G, Dufresne L, Almgren P, Owens DS, Harris TB, Peloso GM, Kerr KF, Wong Q, Smith AV, Budoff MJ, Rotter JI, Cupples LA, Rich S, Kathiresan S, Orho-Melander M, Gudnason V, O’Donnell CJ, Post WS, Thanassoulis G, for the Cohorts for Heart and Aging Research in Genetic Epidemiology (CHARGE) Extracoronary Calcium Working Group. Association of Low-Density Lipoprotein Cholesterol–Related Genetic Variants With Aortic Valve Calcium and Incident Aortic Stenosis. JAMA 2014;312:1764–1771.

48 Del Greco F, Minelli C, Sheehan NU, Thompson JR. Detecting pleiotropy in Mendelian randomisation studies with summary data and a continuous outcome. Statistics in Medicine 2015; DOI: 10.1002/sim.6522

49 Bulik-Sullivan B, Finucane HK, Anttila V, Gusev A, Day FR, Perry JR, Patterson N, Robinson E, Daly MJ, Price AL, et al. An Atlas of Genetic Correlations across Human Disease and Traits. BioRxiv 2015:014498.

50 Law MR, Morris JK, Wald NJ. Use of blood pressure lowering drugs in the prevention of cardiovascular disease: meta-analysis of 147 randomised trials in the context of expectations from prospective epidemiological studies. BMJ 2009; 338:b1665

51 Timpson N, Harbord R, Davey Smith G, Zacho J, TybaergHansen A, Nordestgaard BG. Does greater adiposity increase blood pressure and hypertension risk? Mendelian randomization using Fto/Mc4r genotype. Hypertension 2009;54:84–90

52 Davey Smith G. Use of genetic markers and gene-diet interactions for interrogating level causal influences of diet on health. Genes & Nutrition 2011;6:27–43.

53 Pickrell JK, Berisa T, Segurel L, Tung JY Hinds D. Detection and interpretation of shared genetic influences on 40 human traits. BioRxiv 2015: doi.org/10.1101/019885.

54 The Interleukin 1 Genetics Consortium. IL1 consortium. Cardiometabolic effects of genetic upregulation of the interleukin 1 receptor antagonist : a Mendelian randomisation analysis. Lancet Diabetes Endocrinol 2015; 3: 243–53.

55 Swerdlow DI, Preiss D, Kuchenbaecker KB, Holmes MV, Engmann JE, et al. HMG-coenzyme A reductase inhibition, type 2 diabetes, and bodyweight: evidence from genetic analysis and randomised trials. Lancet 2015;385:351–61.

56 Kathiresan S. Developing medicines that mimic the natural successes of the human genome: lessons from NPC1L1, HMGCR, PCSK9, APOC3, and CETP. J Am Coll Cardiol. 2015 Apr 21;65(15):1562–6.

57 Catapano AL, Ference BA. IMPROVE-IT and genetics reaffirm the causal role of LDL in Cardiovascular Disease. Atherosclerosis 2015;241:498–501.

58 Davey Smith G, Ebrahim S. Mendelian randomization: prospects, potentials and limitations. Int J Epidemiol 2004:33:30–42

59 Davey Smith G. Assessing intrauterine influences on offspring health outcomes: can epidemiological findings yield robust results? Basic and Clinical Pharmacology and Toxicology 2008;102:245–256.

60 Richmond RC, Al-Amin A, Davey Smith G, Relton CL. Approaches for drawing causal inferences from epidemiological birth cohorts: A review. Early Human Development 2014;90:769–780.

61 Lipton P. Inference to the Best Explanation. (2nd edition) Routledge 2004..

